## Supplemental figures for "Human cytomegalovirus evades ZAP detection by suppressing CpG dinucleotides in the major immediate early genes"

#### Supplemental Figure 1

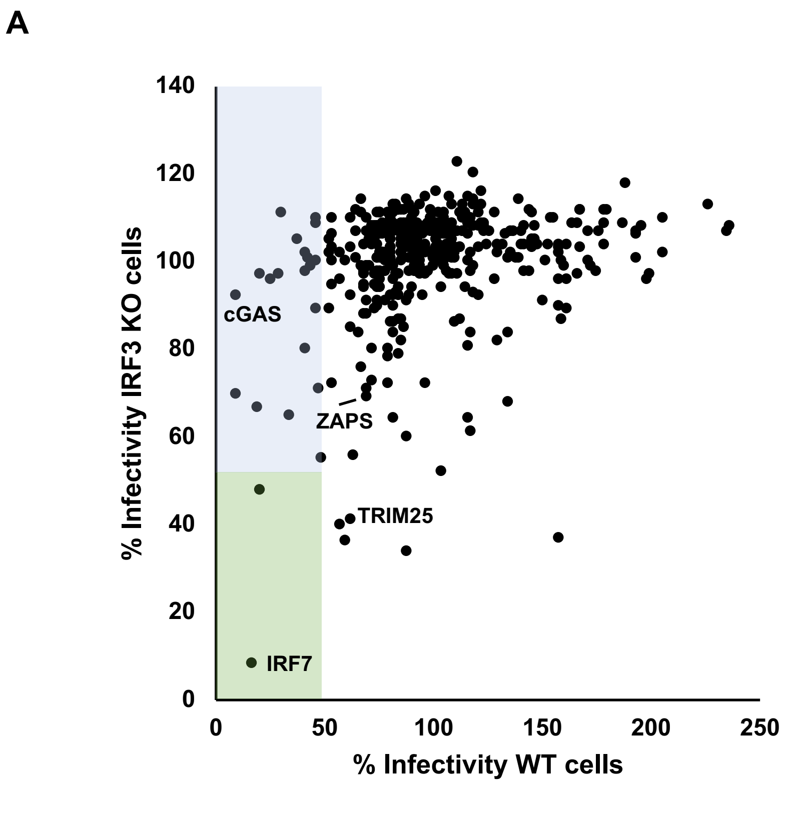


**Supplemental Figure 1.** (A) Direct comparison of relative HCMV primary replication for each individual ISG between wild type and IRF3 KO fibroblast cells. Blue box = ISGs that reduced virus production by more than 2-fold in wild type cells, but were not in IRF3 KO cells. The green box = ISGs that reduced virus production by more than 2-fold in both wild type and IRF3 KO cells. Relative primary replication and virus production levels of HCMV for the ISGs in the blue box (B) and for ISGs that reduced primary replication by more than 50% (C) were plotted.

#### Supplemental Figure 2

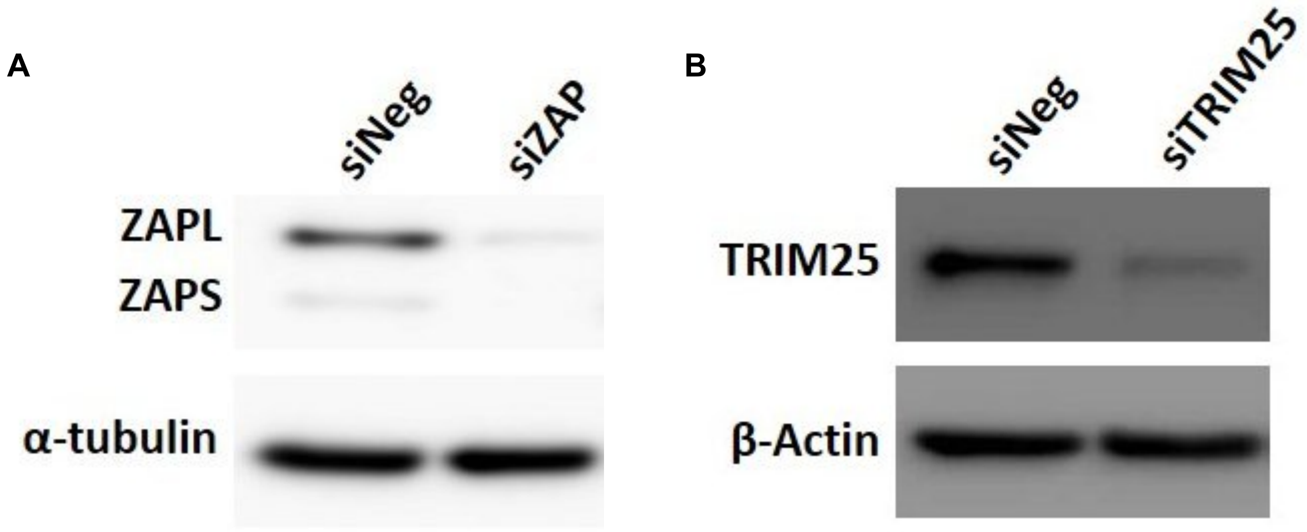


**Supplemental Figure 2. Efficient knockdown of ZAP and TRIM25 by siRNA.** Fibroblast cells were transfected with siRNA targeting ZAP, TRIM25 or a negative control siRNA. Total protein was harvested two days post transfection and levels of ZAP and TRIM25 determined by Western blot analysis.

### Supplemental Figure 3

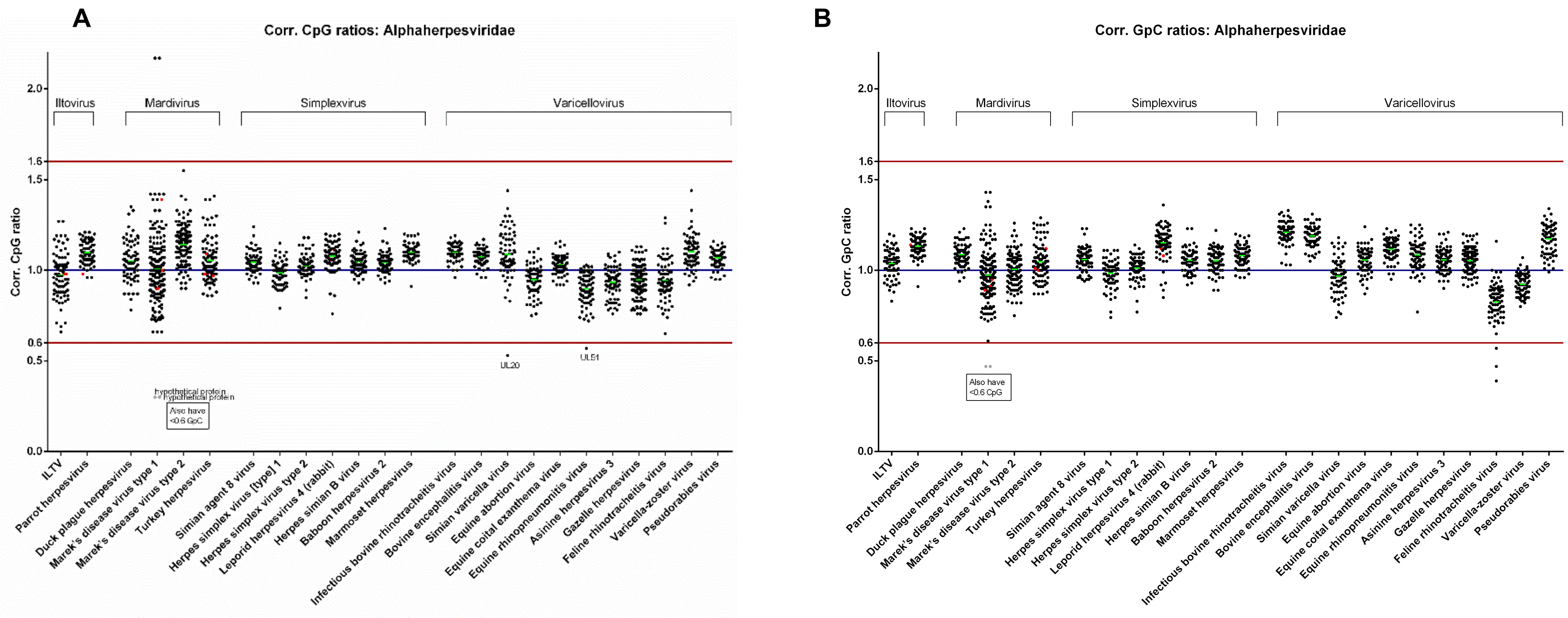


**Supplemental Figure 3. CpG and GpC corrected ratios for ORFs of alpha-herpesviruses.** Corrected CpG and GpC ratios were calculated for alpha-herpesviruses as described in the materials and methods and in figure 3.

#### Supplemental Figure 4

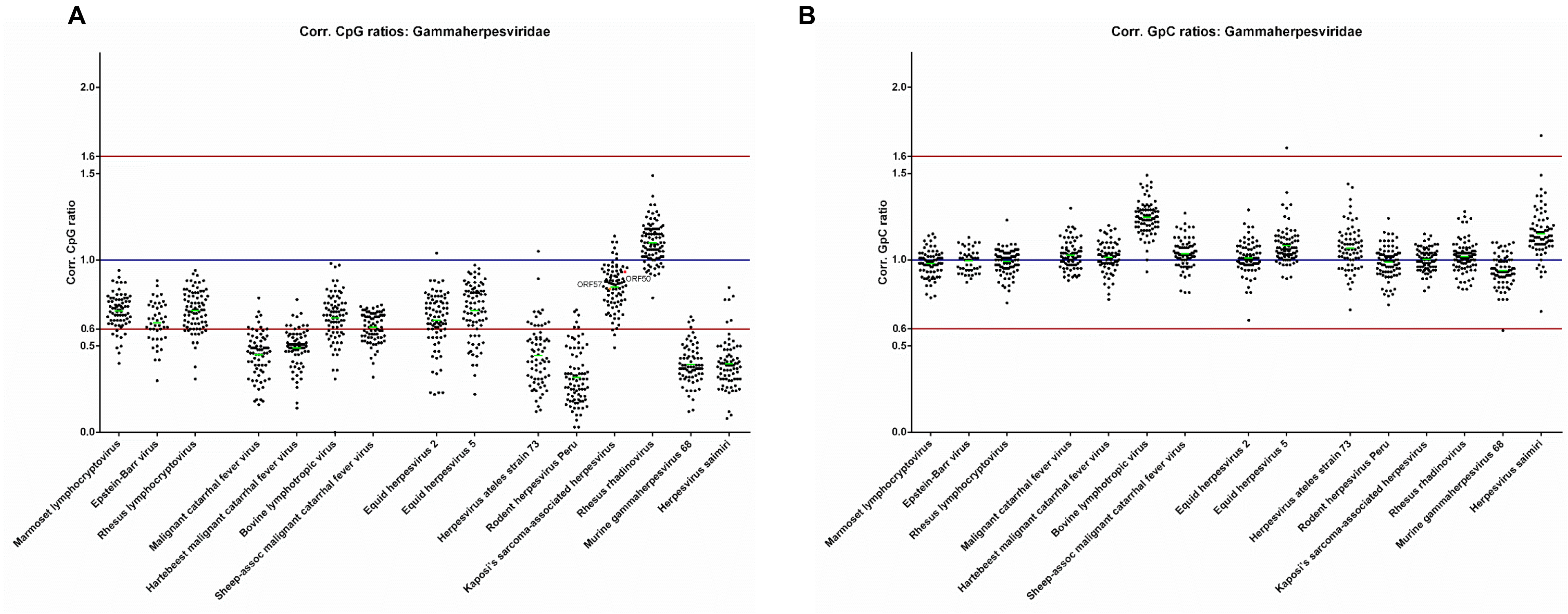


**Supplemental Figure 4. CpG and GpC corrected ratios for ORFs of gamma-herpesviruses.** Corrected CpG and GpC ratios were calculated for gamma-herpesviruses as described in the materials and methods and in figure 3.

#### Supplemental Figure 5

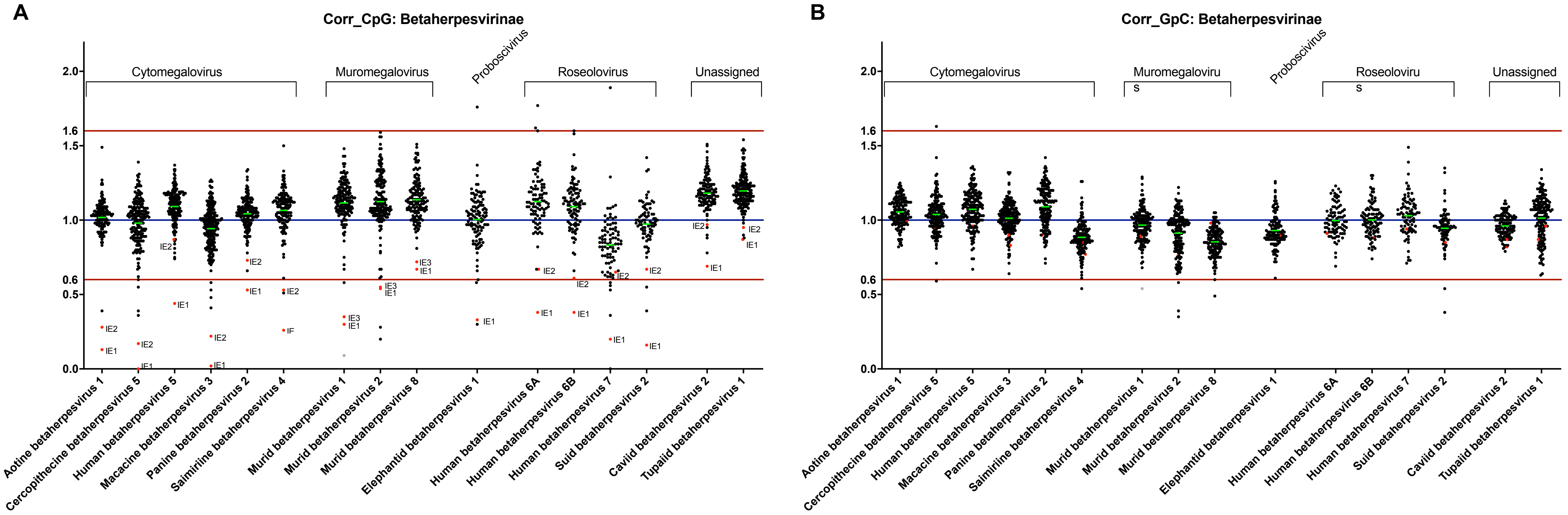


**Supplemental Figure 5. CpG and GpC corrected ratios for ORFs of beta-herpesviruses.** Corrected CpG and GpC ratios were calculated for beta-herpesviruses as described in the materials and methods and in figure 3.

#### Supplemental Figure 6

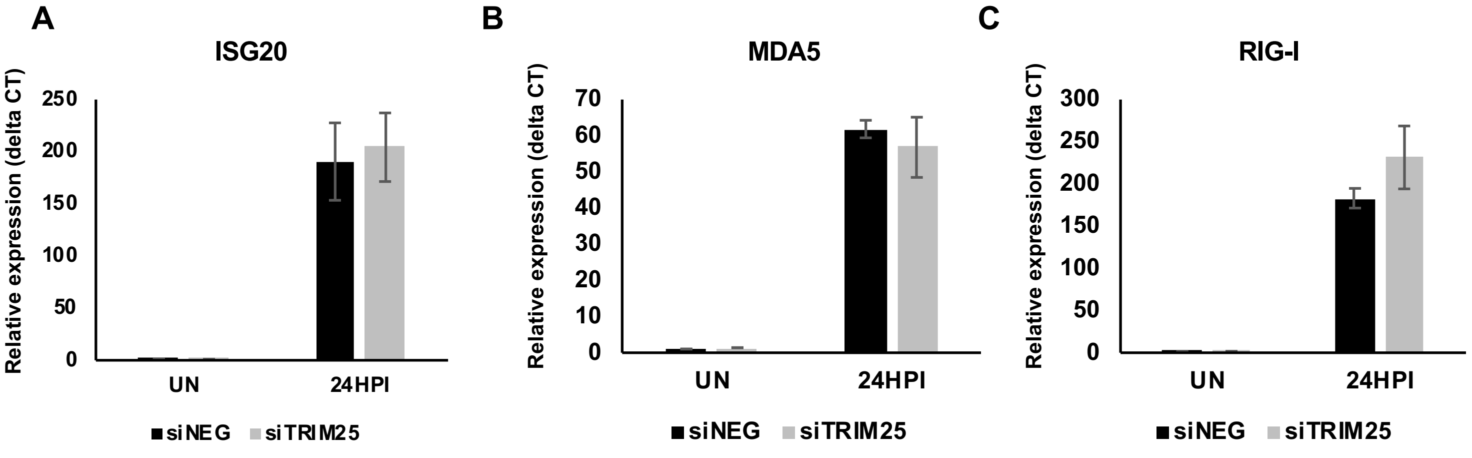


**Supplemental Figure 6. TRIM25 knockdown does not inhibit ISG induction.** Wild-type fibroblast cells were transfected with TRIM25 siRNA or negative control siRNA. 48 hours later, the cells were then infected with TB40E-GFP at an MOI of 3. Total RNA were harvested at 24 hours post infection and the RNA levels of (A) ISG20, (B) MDA5 and (C) RIG-I were determined by quantitative RT-PCR analysis. The result demonstrates that ISG induction following HCMV infection were not inhibited by TRIM25 knockdown.
